# A multi-year survey of helminths from the gastrointestinal tract of wild saddleback (*Leontocebus weddelli*) and emperor (*Saguinus imperator*) tamarins

**DOI:** 10.1101/459099

**Authors:** Gideon A. Erkenswick, Mrinalini Watsa, Alfonso S. Gozalo, Shay Dudaie, Lindsey Bailey, Kudakwashe S. Muranda, Alaa Kuziez, Patricia G. Parker

## Abstract

Noninvasive monitoring of gastrointestinal parasites from wild primates demonstrates that parasite-host relationships are altered during habitat or climatic disturbances. Interpreting changes in parasite measures for population health monitoring is problematic, since wild primates are infected with multiple parasites that fluctuate temporally and seasonally. Individual parasite infection data from two wild populations of New World primates, the saddleback (*Leontocebus weddelli*) and emperor (*Saguinus imperator*) tamarin, were collected over three years to: 1) establish baseline levels of parasite species richness (PSR) and variation across demography; 2) test for non-random associations of parasite co-occurrence; and 3) test hypothesized relationships between group size and PSR. Ten distinguishable parasite taxa were identified from 288 fecal samples by light microscopy following centrifugation and ethyl-acetate sedimentation. These samples represented 105 unique individuals (71 saddleback and 34 emperor tamarins), across 13 saddleback and seven emperor groups. Of the parasites identified in this study, none were confirmed as host specific, and only two parasites had statistically different prevalence between the host species. With few exceptions, individual infection status remained relatively unchanged over the study period. Considering yearly pair-wise parasite associations, we detected no marked differences between expected and observed levels of co-infection, nor did we detect statistically significant associations between group size and parasite species richness over 30 group-years. Logistic models of individual infection status did not identify a sex bias; however, age or species predicted the presence of four and three parasite taxa, respectively. Our model found higher PSR for saddleback tamarins. Considering the two most common parasites, one is typically pathogenic and the other is not, reinforcing caution when translating clinical findings of pathology to real-world systems. We now have reliable baseline data for future monitoring of these populations. Next steps should involve the molecular characterization of these parasites, and the exploration of linkages with health parameters.

## Introduction

Parasitism has a fundamental role to play in the persistence of animal populations in nature, and the richness of parasite communities may serve as effective population- and ecosystem-level measures of health (Hudson, 1998; Hudson, Dobson, & Lafferty, 2006). Evidence shows that natural infections with multiple parasites in wildlife and human populations can have positive effects on host health (Petney & Andrews, 1998). For example, helminth infections have been shown to reduce allergic, autoimmune and inflammatory reactions (Maizels & McSorley, 2016), and helminth-modulated macrophages are now being studied as possible therapies for inflammatory diseases, such as diabetes, multiple sclerosis, and bowel disease (Steinfelder, O’Regan, & Hartmann, 2016). Thus, while “parasitism” connotes the acquisition of resources (space, food, etc.) by one organism at the expense of another, we should be cautious about considering parasites as detrimental in complex environments where they may play a role in maintaining an ecological balance that is necessary for species persistence.

Gastrointestinal parasites, which can be detected through noninvasive surveying, may be ideally suited for health monitoring efforts of a primate community (Gillespie, 2006; Howells, Pruetz, & Gillespie, 2011). They are relatively easy to evaluate from fecal samples collected from habituated primate groups, and can also be acquired in the absence of habituation by scat detection dogs or by searching beneath known feeding or resting locations (Arandjelovic et al., 2015; Orkin, Yang, Yang, Yu, & Jiang, 2016). Several long-term research programs have successfully used temporal parasite data to examine ecological perturbations of threatened primate populations (Bakuza & Nkwengulila, 2009; Chapman, Gillespie, & Speirs, 2005; Gillespie & Chapman, 2008; Gillespie, Chapman, & Greiner, 2005). In contrast, in the absence of temporal data, comparative studies between isolated and more urban primate populations are effective at evaluating impacts of increased contact with humans (Salzer, Deutsch, Raño, Kuhlenschmidt, & Gillespie, 2010; Wenz, Heymann, Petney, & Taraschewski, 2009). Despite the utility of such studies, parts of the world with the highest primate diversity, such as the Neotropics, remain inadequately sampled for naturally occurring gastrointestinal parasites (reviewed in Hopkins and Nunn (2007).

Except for a handful of South American primate taxa, most notably howler monkeys (*Alouatta* spp.), golden lion tamarins (*Leontopithecus rosalia*), and golden-headed lion tamarins (*Leontopithecus chrysomelas*) (Milton, 1996; Monteiro, Dietz, Beck, et al., 2007a; Monteiro, Dietz, Raboy, et al., 2007b; Stuart et al., 1998; Valdespino, Rico-Hernández, & Mandujano, 2010), patterns of parasite-host relationships have been tested primarily among primates in Africa and Asia. These works highlight that parasitism varies with host population demographic variables, including age class and sex (Clough, Heistermann, & Kappeler, 2010; Gillespie, Barelli, & Heistermann, 2013; Gillespie et al., 2010; Macintosh, Hernandez, & Huffman, 2010), and sexual maturity or dominance (Macintosh et al., 2012; Muehlenbein & Watts, 2010), although not all studies concur (Gillespie et al., 2010; Setchell et al., 2006; Setchell, Charpentier, & Abbott, 2009). Host behavior in combination with parasite mode of dispersal can also structure parasite populations (Macintosh et al., 2010; Nunn & Heymann, 2005). Influences of concomitant parasite infections are not routinely analyzed, but when they are, their impacts may be comparable to those exerted by host or environmental factors (Erkenswick, Watsa, Gozalo, Dmytryk, & Parker, 2017a; Monteiro, Dietz, Raboy, et al., 2007b; Nunn, Brezine, Jolles, & Ezenwa, 2014; Telfer et al., 2008). Also, meta-analyses find general support for increasing parasite species richness as social group size increases (Cote & Poulin, 1995; Nunn, Altizer, Jones, & Sechrest, 2003; Rifkin, Nunn, & Garamszegi, 2012; Vitone, Altizer, & Nunn, 2004), but the spatial and taxonomic scales at which this pattern will hold true (e.g. within a single species or population) require further study. Finally, parasite populations may also vary across seasons and years (Clough et al., 2010; Gillespie et al., 2010).

Since the range of factors that can explain parasite-host patterns in nature is large, often dependent on the environment and time, it may be best approached through longitudinal monitoring of individuals in host communities (Clutton-Brock & Sheldon, 2010; Stuart et al., 1998). A primary challenge has been that research on wild primates requires habituation to observers, which often constrains sample sizes, making it difficult to analyze many of the factors (Williamson & Feistner, 2011). Thus far many studies have offered snapshots of parasite prevalences, focused on just one or two parasites of known interest, the sampling of a single primate host, or on data from health inspections, or necropsies, after animal extraction from the wild (Cosgrove, Nelson, & Gengozian, 1968; Porter, 1972; Wolff, 1990). Collectively they have created a broad foundation of primate parasite data (see Nunn & Altizer, 2005 for a detailed compilation). Emerging patterns can now be examined carefully in the wild to look at influences of host demography and development, mode of transmission, and change over time at the level of a population. As an example, for almost a half-century it has been well known that New World monkeys are broadly infected by *Plasmodium brasilianum*, a quartan malarial parasite, that may in fact be the same as the human parasite *Plasmodium malariae* (Collins & Jeffery, 2007; Lalremruata et al., 2015). However, only last year do we have the first evidence that it may persist in a highly aggregated manner among a small number of chronically infected non-human primate hosts (Erkenswick, Watsa, Pacheco, Escalante, & Parker, 2017b). In addition, long-term studies that incorporate more than one primate host individual are essential to examine several longstanding hypotheses of how sociality influences parasite prevalence, intensity, and diversity (Altizer et al., 2003; Freeland, 1976; 1979), as are long-term studies of multiple sympatric species to examine species-specificity of infection dynamics.

The Callitrichidae (comprised of tamarins and marmosets) are small arboreal primates that are widely distributed throughout the forests of South America (Sussman & Kinzey, 1984). They are frequently found in sympatry with other New World monkeys and in some cases have proven relatively resilient and flexible in the face of encroachment by human populations (Gordo, Calleia, Vasconcelos, Leite, & Ferrari, 2013; G. C. Leite, Duarte, & Young, 2011; Soto-Calderon, Acevedo-Garces, Alvarez-Cardona, Hernández-Castro, & Gartia-Montoya, 2016). Part of their ecological flexibility may be due to their generalist diets that include fruits, insects, tree exudates, and fungi (Sussman & Kinzey, 1984), a characteristic that also could expose them to a wide array of parasites that are dispersed by intermediate arthropod hosts. Studies of the gastrointestinal parasites of callitrichids have documented overlap with other primate families including the Ateledae, Cebidae, and Aotidae (Michaud, Tantalean, Ique, Montoya, & Gozalo, 2003; Phillips, Haas, Grafton, & Yrivarren, 2004; Tantalean, Gozalo, & Montoya, 1990; Wolff, 1990). Considering the approximately 60 species and subspecies of Callitrichidae, there have been only a handful of comprehensive evaluations of gastrointestinal parasites from free-ranging populations (Monteiro, Dietz, Beck, et al., 2007a; Müller, 2007; Wenz et al., 2009), and only two species in which parasites have been monitored routinely over time, golden lion and golden-headed lion tamarins (*Leontopithecus rosalia* and *L. chrysomelas*, respectively (Monteiro, Dietz, Raboy, et al., 2007b).

The principle aim of this study was to characterize the gastrointestinal helminth assemblages from two populations of sympatric, individually identifiable, free-ranging callitrichids - the saddleback tamarin (*Leontocebus weddelli*, formerly *Saguinus fuscicollis weddelli*) (Buckner, Lynch Alfaro, Rylands, & Alfaro, 2015; Matauschek, Roos, & Heymann, 2011) and emperor tamarin (*Saguinus imperator*) - from fecal samples collected noninvasively and via an annual mark-recapture program. Both hosts exhibited group sizes (3 – 8 individuals), mating systems, and reproductive behaviors characteristic of most callitrichids (Watsa, Erkenswick, & Robakis, 2017), where a single, reproductively dominant female mates with multiple males and gives birth to twin offspring once a year, cared for by other adult group members (Sussman & Kinzey, 1984; Wislocki, 1939. By sampling these hosts across three years, we estimated the prevalence of gastrointestinal parasites, parasite species richness, and the extent of parasite overlap between the two host species. We also calculated rates of change in infection status from animals that were screened for helminths in two consecutive years. In doing so, we established baseline data for future comparative studies following perturbations such as changing weather patterns due to climate change, habitat loss/modification, or greater human encroachment. Second, we analyzed how parasite prevalence varies by host demography, age class and sex, and co-infection. As a result of greater social burdens placed on females to compete for dominant breeding opportunities, we predicted that an age-sex interaction would influence prevalence and parasite species richness. Specifically, we predicted that adult females of both host species would have higher prevalence and richness. Based on prior research of blood parasites from these populations (Erkenswick et al., 2017a), we predicted that there would be non-random prevalence of several co-infections, considering all pairwise combinations of parasites. Finally, we tested the hypothesis that there would be a relationship between group size and parasite species richness, and predicted that larger groups would harbor greater numbers of parasites, which has not yet been tested within the Callitrichidae. Our findings are discussed in terms of parasite pathogenicity and parasite mode of dispersal.

## Methods

### Field site and study subjects

Sample collection took place annually from 2012 - 2014 in the Madre de Dios Department of Southeastern Perú at the Estacion Biologica Rio Los Amigos (EBLA) (12°34’07”S, 70°05’57”W), which is managed by the Asociacion para la Conservation de La Cuenca Amazonica. All sampling took place within a forest trail system that covers approximately 900 ha of tropical rainforest that is adjacent to the Los Amigos Conservation Concession inside the buffer zone of Manu National Park. There are two distinct seasons each year at this site – the wet season from October to March, (average monthly precipitation > 250 mm), and the dry season from April to September (136 mm ± SD 19 mm) (Watsa, 2013). All sampling took place during the dry season, from May – July each year, precluding the study of the effects of seasonality on the parasite community in these primates.

Three callitrichines at this site, the saddleback tamarin, emperor tamarins, and the more cryptic Goeldi’s monkey (*Callimico goeldii*) (Watsa, Erkenswick, Rehg, & Pitman, 2012), share forest habitat with eight other primate species including three species of Cebidae, and two species each of Atelidae and Pithecidae, as well as owl monkeys (*Aotus nigrifrons*) (Watsa, 2013). At EBLA, both *S. imperator* and *L. weddelli* have average group sizes of 5 (range of 3-8) individuals and group compositions are similar (Watsa et al., 2017; 2015). The primary differences between *S. imperator* and *L. weddelli* are adult weight, 515g ± 66 and 386g ± 86, respectively, and nuances in feeding behavior including greater amounts of fungi consumption in *S. imperator* (pers. obs.).

Each individual sampled was classified into one of three age classes based on dental eruption patterns (Watsa, 2013). Juveniles were defined as individuals whose adult teeth were absent or not fully erupted (<11 months old). Sub-adults were animals with adult teeth, but that were juveniles in the preceding year. All remaining individuals were assigned to the adult age class. Due to small sample sizes from the sub-adult class, the juveniles and sub-adult classes were combined to analyze the effects of age on parasite prevalence.

### Sample collection and storage

Since 2009, an annual mark-recapture program has been implemented on ~ 70 saddleback and emperor tamarins by Field Projects International (Watsa et al., 2015). During capture, each individual is permanently tagged with a Home Again microchip, and was made visually identifiable by unique patterns of bleached rings around the tail, as well as a tricolor beaded necklace that signified group, sex and individual identity (for the full capture protocol see Watsa et al., 2015). In addition to collecting fecal samples at the time of capture, we used radio telemetry to track tamarins in 14 groups each year via a radio collar placed on the breeding female in each group (Wildlife Materials, Murphysboro IL). We also used both full (sleep-site to sleep-site, spanning ~ 11 hours) and half-day (minimum 5 hours) follows to opportunistically collect fecal samples from all group members as they were produced.

Upon collection during mark-recapture and follows, all fecal samples were transferred using sterile technique into numbered plastic bags and stored in a chilled thermos. Upon return to basecamp, each sample was fixed in 10% neutral buffered formalin (1:2, feces to preservative ratio). For each sample, we recorded species, individual ID, group, date, time of day, and type of collection (follow or trapping event). Only samples produced by identified individuals were included in this study. All samples were exported to the Parker Laboratory at the University of Missouri – St. Louis for analyses.

All sampling protocols adhered to guidelines outlined by the American Society of Mammalogists (Sikes & Gannon, 2011) and were approved by the Institutional Animal Care and Use Committee at the University of Missouri-St. Louis and the Directorate of Forest and Wildlife Management (DGFFS) of Perú annually. The DGFFS also granted export permits for the samples, while the CDC and US Fish and Wildlife Services approved the import of these samples into the USA.

### Laboratory analysis

Isolation of parasite cysts, eggs, and larvae from fecal samples followed a two-step process based on sedimentation procedures as per MacIntosh (2010) and Zajac and Conboy (2012). In Step 1, we used a fecal straining procedure in which fecal samples were 1) diluted in 10% neutral buffered formalin, 2) strained of large debris through cheese cloth into a plastic cup, 3) transferred to a 15ml falcon tube with an empty weight already recorded, 4) centrifuged at 800xg for 5 minutes to form a fecal pellet, 5) removed of the supernatant and weighed, 6) re-suspended and homogenized in 5ml of 10% formalin. In Step 2, we followed the centrifugal sedimentation test outlined by Zajac and Conboy (2012) with 1ml of the homogenized suspension from Step 1. Sedimentations from Step 2 were re-suspended in exactly 1ml of preservative, and 80ul aliquots were placed onto clean slides with coverslips for full evaluation with an Olympus CX31 light microscope using 200x magnification (micrographs were taken at higher power). Evaluations of parasites were timed and tabulated using a free online data counter, COUNT (http://erktime.github.io/count/), and each unique infection/sample was documented with multiple micrographs taken with a Leica ICC50 HD camera. Three separate aliquots per sample were evaluated with each evaluation taking an average of 10 minutes.

Unless infections were too rare, standard length and width measurements from 10 representative micrographs per parasite per species were recorded with a calibrated ruler in Image J (https://imagej.nih.gov/ij/) to the nearest 1 μm. Measurements of all parasite forms were compared to known references values in the literature and identified to the lowest taxonomic scale possible.

### Statistical analysis

Average prevalence, as well as the proportion of individuals that acquired infection, lost infection, or showed no change in infection status, was calculated for each helminth identified by microscopy across the three-year study period. Average change in infection status was determined by selecting all instances where an individual was sampled across a two-year period, either 2012-2013 or 2013-2014, and computing the mean number of individuals that acquired, lost, or did not change infection status. Differences in helminth prevalence between host species were tested with Fisher’s Exact Test and adjusted p-values following the Holm-Bonferroni method (Holm, 1979). To test for variation in the presence of parasitic infections across host variables we used mixed-effect logistic regression models with a binary response variable and binomial errors. Fixed effects included ‘sex’, ‘age class’, and ‘species’ and random effects included ‘animal identity’ and ‘year’ to accommodate individual resampling and possible inter-annual variation. We also incorporated the number of samples collected per animal per year as an offset to account for temporal sampling bias (Walther, Cotgreave, Price, & Gregory, 1995). Parasite species richness, which was a discrete numerical response variable, was analyzed with an identical model formula but using Poisson errors. Model selection for all models was carried out with step-wise term deletion by removing non-significant factors and comparing nested models with a likelihood ratio test.

To test for significant correlations between group size and parasite species richness we calculated ratified parasite community richness estimates per group. The use of species accumulation curve estimates are advocated by Walther et al. (1995), because raw values of parasite community richness are easily biased by uneven sampling. We used Spearman’s rank correlations to test if parasite community richness estimates with similar sampling effort were associated with group size.

To identify any nonrandom parasite co-occurrences, we compared the prevalence of all observed pairwise co-infections with expected estimates of co-infection (calculated as prevalence of A * prevalence of B). We then plotted expected against observed values to identify discordant levels of co-infection, and if applicable, used a two-sample z-test to compare proportions. All statistical analyses were performed in R (R Development Core Team, 2015).

## Results

In total, we collected 288 individually identified fecal samples from 105 unique tamarins (71 *L. weddelli*, 34 *S. imperator*) distributed across 13 groups of *L. weddelli* and 7 groups of *S. imperator.* The number of samples collected per individual per year ranged from 1 to 7, with a mean of 1.6, and median and mode of 1. The average fecal sample weight, following Step 1 in sample processing (see Methods), was 0.41grams +/− 0.22. Considering all sex and age classes, our sampling included slightly more males than females across years, and sub-adults of both hosts species were the least sampled age group (Table 1).

**Table 1.**
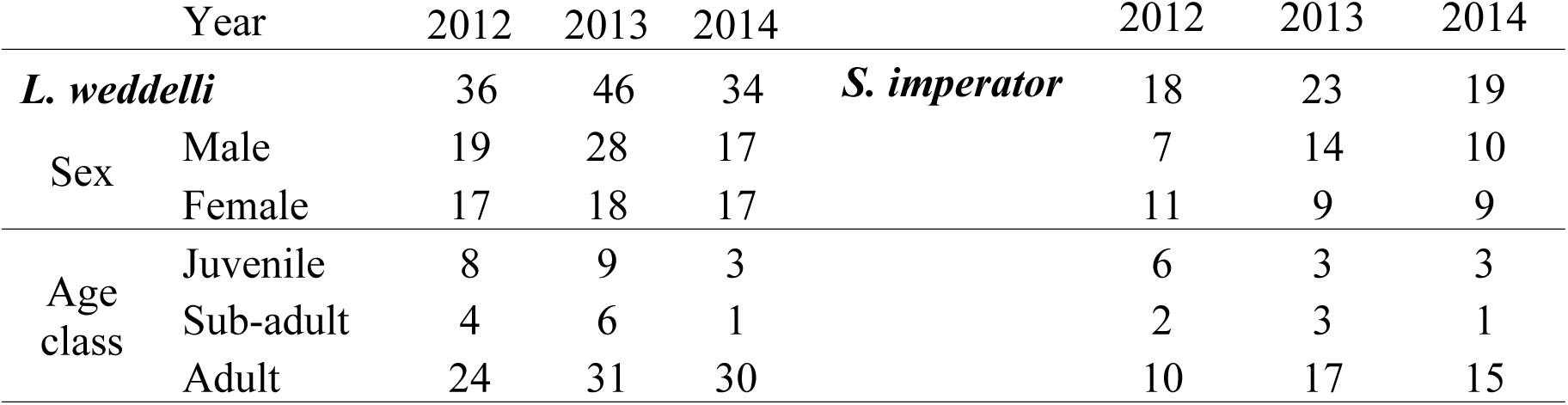
Numbers of individuals sampled by species, sex, age class, and year

| Year |  | 2012 | 2013 | 2014 | Year |  | 2012 | 2013 | 2014 |
| --- | --- | --- | --- | --- | --- | --- | --- | --- | --- |
| ***L. weddelli*** |  | 36 | 46 | 34 | ***S. imperator*** |  | 18 | 23 | 19 |
| Sex | Male | 19 | 28 | 17 |  |  | 7 | 14 | 10 |
|  | Female | 17 | 18 | 17 |  |  | 11 | 9 | 9 |
| Age class | Juvenile | 8 | 9 | 3 |  |  | 6 | 3 | 3 |
|  | Sub-adult | 4 | 6 | 1 |  |  | 2 | 3 | 1 |
|  | Adult | 24 | 31 | 30 |  |  | 10 | 17 | 15 |

We were able to differentiate 10 helminth parasites by morphology. We identified 8 to a likely species or genus, and 2 to the level of family. Most of these parasites have been detected in the Callitrichidae in the past (See Supplemental Fig. 3 for representative micrographs, standard measurements, and additional citations). All but one rare parasitic infection, *Spirura guianensis*, were found in both host species, although prevalence profiles varied (Table 2). Prevalence for the Dicrocoeliidae was significantly higher in *S. imperator* (Fisher’s test mean-adjusted P-values = 0.012), and Cestoda was significantly higher in *L. weddelli* (Fisher’s test mean adjusted P-value = 0.008).

**Table 2.**
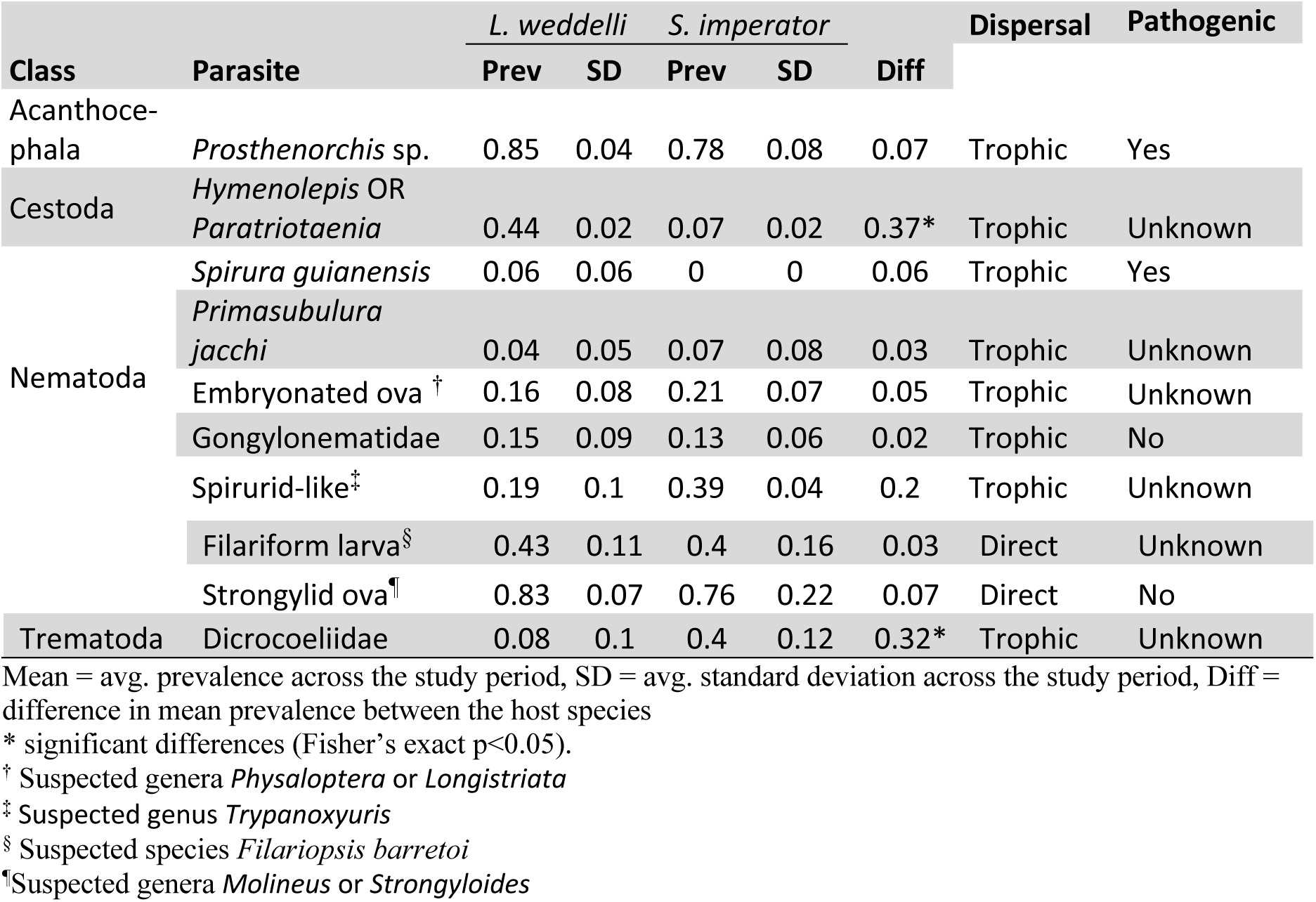
Mean annual prevalence by host species and parasite

For most helminths, individual infections remained unchanged with small proportions of individuals switching status in both directions (Fig. 1). Among *S. imperator*, this pattern differed for Strongylid ova and Dicroceoliidae (Fig. 1, Supplemental Table 4). Among *L. weddelli*, we only detected frequent changes in individual infection status for Cestoda. (Fig. 1, Supp. Table 4). Our rarest parasite infections for both host species were *Spirura guianensi* and *Primasubulura jacchi*, but Cestoda for *S. imperator* in particular.

**Figure 1.**
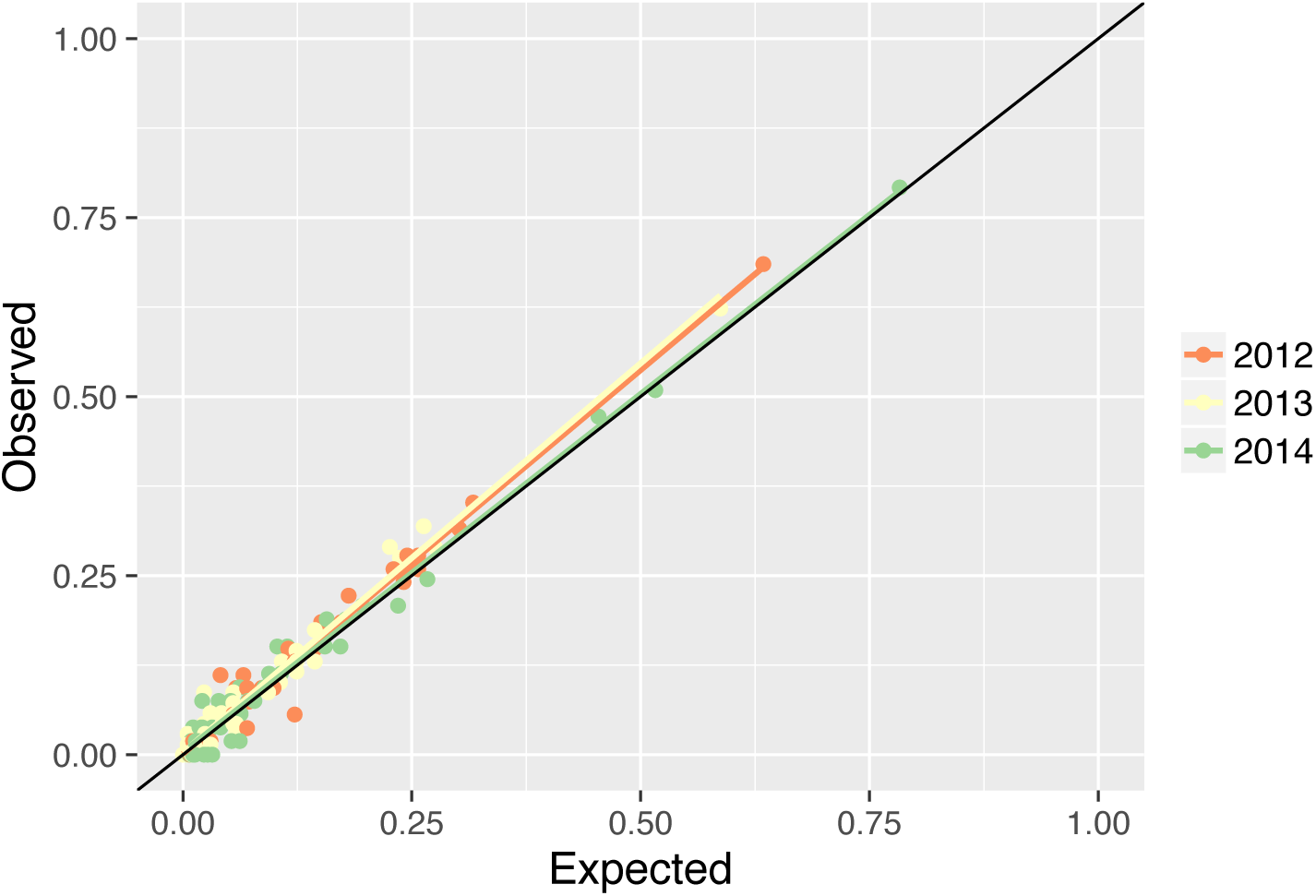
Observed versus expected prevalence of parasite co-infection. Each dot represents a unique pairwise combination of parasites.

Considering each year separately, we did not detect any significant deviations between expected and observed prevalence of co-infection (Fig. 2); the largest absolute difference in prevalence across all parasite combinations throughout the study period was 0.07. We also found no evidence of a relationship between group size (ranged 3 – 8) and estimated parasite species richness within groups after controlling for sampling effort (Spearman’s rank correlation = −0.08, P-value = 0.665, n = 30).

**Figure 2.**
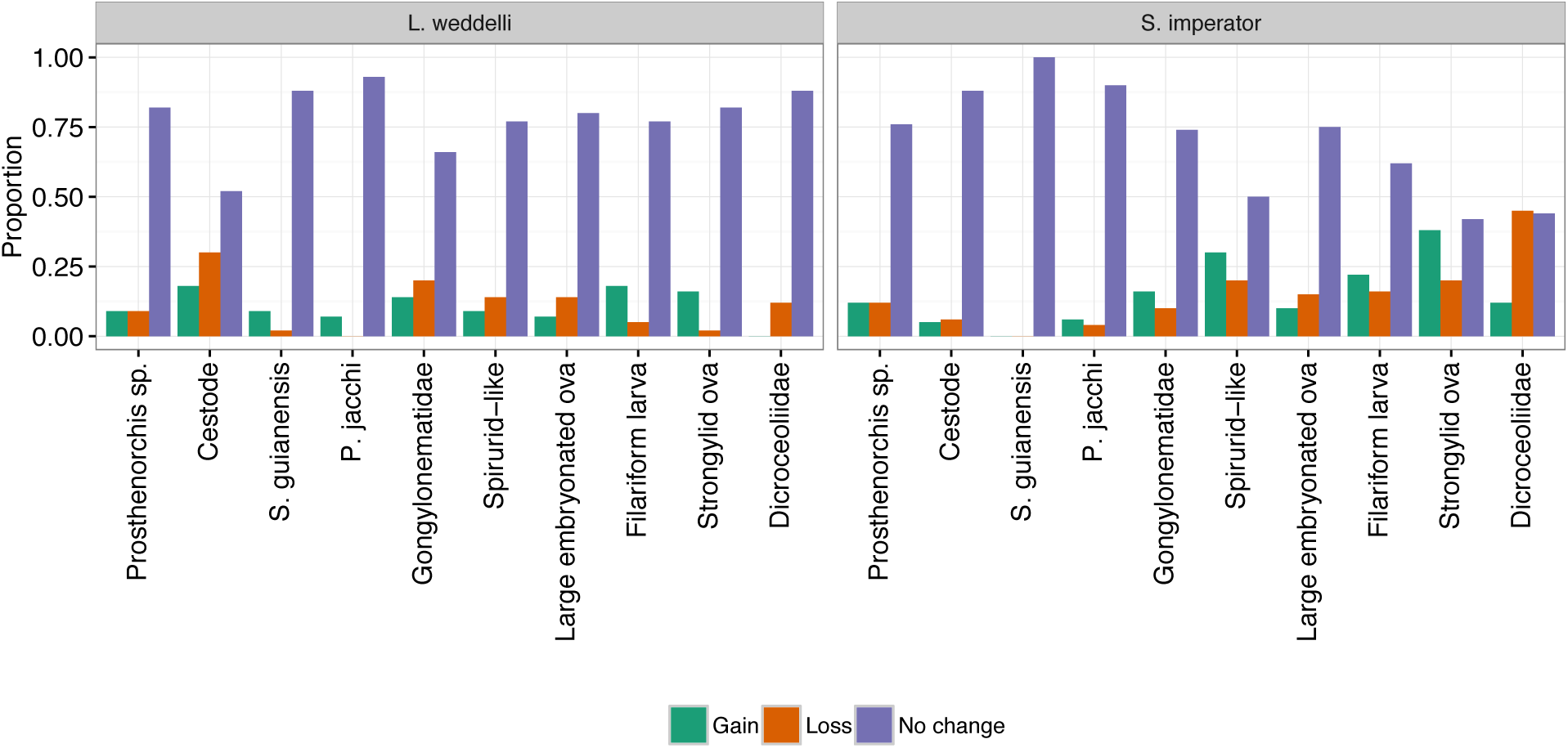
Average proportion of individual infection status change by host species and parasite.

Out of the 10 helminths identified, 6 were common enough to evaluate their distributions across host species, sex and age class variables; *Spirura guianensis*, Gongylonematidae, and *Primasubulura jachii* were too rare to analyze prevalence patterns using statistical models. No parasitic infection exhibited a significant sex bias; however, age and species did predict the presence of 4 and 3 parasites, respectively (Table 3). Relative to *L. weddelli, S. imperator* was positively associated with Dicrocoeliidae but negatively associated with cestode and strongylid ova. Relative to adults, juveniles and sub-adults were negatively associated with *Prosthenorchis* and filariform larva, but positively associated with cestode and the large embryonated ova. Our models of parasite species richness identified host species as the only significant predictor considered in this study (Table 3), which had a significantly negative estimate for *S. imperator.*

**Table 3.**
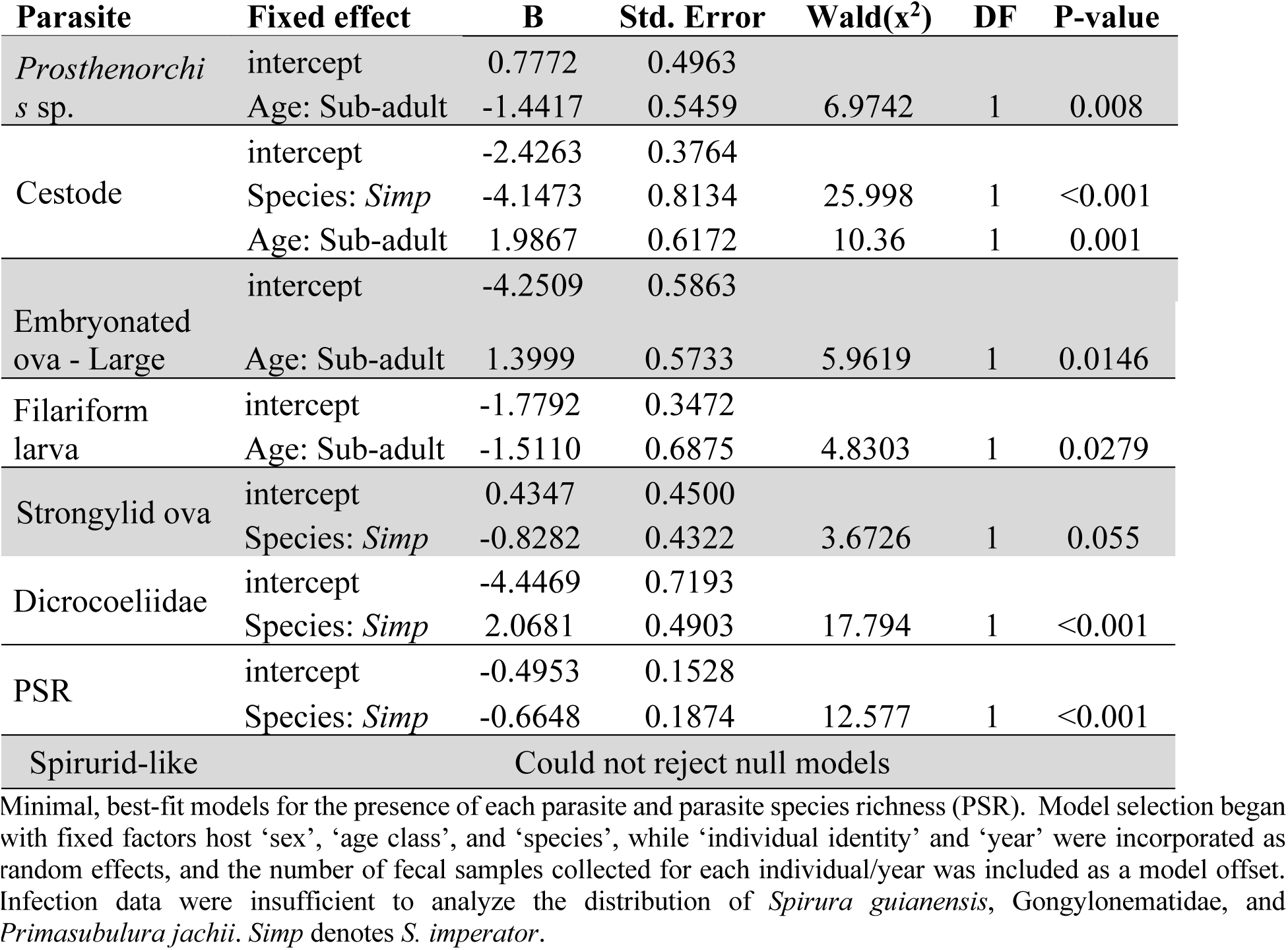
GLMM outcomes for each parasite and parasite species richness

## Discussion

It is customary for wild animals to acquire and maintain multiple parasitic infections during their lives (Cox, 2001; Petney & Andrews, 1998). While clinical and/or experimental studies are effective at demonstrating the pathogenicity of parasites, translating such findings to natural systems is not always straightforward. For example, the thorny-headed worms (Phylum Acanthocephala) are well-known parasites of Central and South American primates (King, 1993; Tantalean et al., 1990). They attach themselves to the intestinal mucosa of their primate hosts and cause inflammatory responses, obstruction of the lumen, and the formation of lesions and ulcers, which may lead to secondary infections or even peritonitis in the worst cases (King, 1993; Strait, Else, & Eberhard, 2012). In spite of the potential to be extremely pathogenic, Acanthocephala infections (e.g. *Prosthenorchis* spp.) are common in this study population, routinely found in other surveys of callitrichids (Müller, 2007; Tantalean et al., 1990; Wenz et al., 2009), and are also present in other New World primate families such as the Cebidae and Atelidae (King, 1993; Phillips et al., 2004; Wenz et al., 2009). It has been shown before that in nature, where the host’s ability to tolerate infection is most important to its fitness, parasite pathogenicity may be dictated by environmental factors (Cardon, Loot, Grenouillet, & Blanchet, 2011) that Walker et al. (2010) discussed as “environmental forcing of pathogenicity”. On top of this, interactions between diverse parasite species modulate pathogenicity (Balmer, Stearns, Schötzau, & Brun, 2009; Lello, Boag, Fenton, Stevenson, & Hudson, 2004; Monteiro, Dietz, Raboy, et al., 2007b; Petney & Andrews, 1998). Of the 9 helminths documented in this study, 5 are of unknown pathogenicity, 2 are probably non-pathogenic, and 2 are known to be pathogenic (Table 2). That one of two most common parasites in this study, across years, is considered highly pathogenic (*Prosthenorchis*), suggests taking caution before transferring clinical findings to real-world systems. In this study population, individuals have been hosting *Prosthenorchis* for upwards of 12 years [Erkenswick, unpublished data], thus observations of particularly pathogenic parasites (e.g. Acanthocephalans) should be viewed in context with the broader parasite community and changes in the environment, which requires detailed longitudinal data collection (Haukisalmi, Henttonen, & Tenora, 1988).

Although our analysis did not identify nonrandom associations between co-infecting parasites, it is still possible that within-host parasite interactions are at play. The use of presence-absence infection data is much less sensitive at detecting relationships than accurate measures of parasite intensity or burden (Knowles et al., 2013; Lello et al., 2004), which will be an aim of future studies. Our inability to detect a relationship between group size and parasite species richness is consistent with our previous work on blood parasites (Erkenswick et al., 2017a), and may be a consequence of too little variation in group sizes, which ranged only from 3 to 8. Other genera of callitrichids can occur in slightly larger groups than our hosts species; for example, *Callithrix* groups can be as large as 15 members (Pontes & da Cruz, 1995; Watsa et al., 2017), but it also may be that generalities about group size and parasite diversity do not apply within the Callitrichidae.

The data we have provided here represent the first description of the intestinal parasite for free-ranging *Saguinus imperator.* Although the IUCN currently lists *S. imperator* of least concern (Rylands & Mittermeier, 2008), the core of its distribution is surrounded by one of the fastest growing gold mining industries in the world (Asner, Llactayo, Tupayachi, & Luna, 2013). Moreover, *S. imperator* is currently one of the most valuable Peruvian monkeys in the illegal wildlife trade (Watsa, 2015). Hence, close monitoring of intact populations of *S. imperator* is crucial to its conservation. We also provide a new comparative dataset of gastrointestinal parasites from *L. weddelli* (formerly *S. fuscicollis weddelli*), which is also a primate of ‘least concern,’ though recent taxonomic revisions may result in revision of its conservation status (Buckner et al., 2015; Matauschek et al., 2011).

We find some noteworthy similarities and differences between this study and previous studies of gastrointestinal parasites from congeneric callitrichids. Phillips et al. (2004) screened one group of *S. fuscicollis* in the nearby Tambopata National Reserve and identified four parasites (*Trichuris*, *Iodamoeba, Entamoeba*, and an unidentified strongyle), none of which were confirmed in our hosts. Although they had a small sample size of 4 individuals, it is surprising that they did not detect *Prosthenorchis* sp. infection, which they did find in 1 of 18 squirrel monkey (*Saimiri sciureus*) fecal samples at the same site. In northern Peru, both Wenz et al. (2009) and Muller (2007) conducted gastrointestinal parasite surveys over a single season from sympatric callitrichine hosts, *S. fuscicollis* and *S. mystax*, and reported a parasite assemblage that mostly overlaps with our findings (*Prosthenorchis, Hymenolepis*, large and small Spirurids, *Primasubulura*, Strongylid larvae). Both recorded higher prevalence than our study for every helminth except *Prosthenorchis*, which was considerably less common. Comparing both of their tamarin hosts, they detected higher prevalence of *Hymenolepis* and PSR for *S. fuscicollis*, as we did for *L. weddelli* (until recently, *S. fuscicollis* and *L. weddelli* were consider the same species (Buckner et al., 2015; Matauschek et al., 2011)). The variation in prevalence between the studies is likely related to ecological differences between the locations. We suspect that greater diversity of helminths observed in this study is associated with a higher diversity of primates (11 compared to 4), but also may be a consequence of prior human activities at the site which altered the densities of primate species at EBLA (Rosin & Swamy, 2013).

Interpreting these findings in light of parasite mode of transmission, direct or trophic, is challenged by the lack of information on the exact intermediate hosts of many parasites, and many could have more than one. The majority of the parasites we detected were trophically transmitted – encountered via an intermediate host (usually an arthropod)– and prevalence significantly differed between host species in two instances. While a comprehensive study of feeding and foraging behavior has not been conducted on our host populations, it has been conducted on sympatric *S. fuscicollis* and *S. mystax* in Northern Perú. The findings from two studies on tamarin feeding ecology agree that *S. fuscicollis* spends significantly more time foraging in the lower strata and on the ground, while the opposite was true of *S. mystax* (Heymann, Knogge, & Tirado Herrera, 2000; Smith, 2000). Smith (2000) documented that sympatric, congeneric host species exercised distinct feeding preferences such as color and size of prey items, and this might account for host differences observed here for Dicrocoeliidae and Hymenolepis infection if prey items are unevenly distributed across forest habitat. Niche specialization might also contribute to the variation we observed in parasite species richness among these hosts via differences in intermediate host encounter rates or the persistence of parasite free-living phases on the ground or in certain forest strata. Consistent with the prevalence of blood parasites from this population (Erkenswick et al., 2017a), age class predicted the presence of two trophically transmitted parasites, though we obtained both positive and negative relationship estimates from our models. We attribute this to differences in diet and foraging efficiency between younger and older individuals, which would alter parasite encounter rates.

By tracking prevalence of parasitic infections over time in wild populations it is possible to infer whether natural parasite communities are stable; however, longitudinal data at the level of the individual provides insights into the source, or lack, of population stability (Knowles et al., 2013). In some cases, it could even aid in identification of parasites that have negative health consequences. For example, if parasite prevalence is consistently low relative to the incidence of new infections across years, and there is little evidence that individuals clear infections, then previously infected hosts must be disappearing regularly. In this study, we encountered few disparities in the rate of acquisition or loss of parasites, and considered in concert with observed prevalence, we see no obvious signs of negative health consequences.

The results provided here, in combination with recent works on hemoparasites (Erkenswick et al., 2017a), represent a benchmark against which future parasitological surveys can be compared. Represented in these callitrichine host parasite assemblages are parasites that are transmitted directly, trophically, or by arthropod vectors. Given changes in the environment that alter food availability or vector populations, we would expect corresponding deviations from what has been documented here. The near ubiquity across South America rainforests, propensity to be found in sympatry with other New World primates, and exist in and around human-altered landscapes, make the Callitrichidae a potential flagship family for the regional detection of ecological changes, or even environmental threats.

## Acknowledgements

We wish to thank research assistants from Field Projects International who aided in the collection of tamarin fecal samples. We want to thank Cindee Rettke for her assistance coordinating wet laboratory analyses, and Aaron Erkenswick for creating a web application that was used to record infection data. We express our gratitude to the following agencies and organizations that provided non-monetary support to this investigation: Centro de Ecologia y Biodiversidad, Amazon Conservation Association, el Estacion Biologica Rio Los Amigos. This work was supported by Field Projects International, the American Society of Mammalogists, University of Missouri Trans World Airlines Scholarship Program, Whitney R. Harris World Ecology Center, Sigma Xi Society, IdeaWild, and the Intramural Research Program of the National Institutes of Health, National Institute of Allergy and Infectious Diseases, Comparative Medicine Branch.

All research protocols and data collection in this paper were reviewed and approved annually by the IACUC of the University of Missouri-St. Louis as well as el Servicio Nacional Forestal y de Fauna Silvestre (SERFOR) of Peru. This research also complied with the American Society of Primatologists Principles for the Ethical Treatment of Non-Human Primates.

## References

Altizer, S., Nunn, C. L., Thrall, P. H., Gittleman, J. L., Antonovics, J., Cunningham, A. A., et al. (2003). Social organization and parasite risk in mammals: integrating theory and empirical studies. Annual Review of Ecology, Evolution and Systematics, 34, 517–547. http://doi.org/10.1146/annurev.ecolsys.34.030102.151725

Arandjelovic, M., Bergl, R. A., Ikfuingei, R., Jameson, C., Parker, M., & Vigilant, L. (2015). Detection dog efficacy for collecting faecal samples from the critically endangered Cross River gorilla *(Gorilla gorilla diehli)* for genetic censusing. Royal Society Open Science, 2(2), 140423–140423. http://doi.org/10.1098/rsos.140423

Asner, G. P., Llactayo, W., Tupayachi, R., & Luna, E. R. (2013). Elevated rates of gold mining in the Amazon revealed through high-resolution monitoring. Proceedings of the National Academy of Sciences of the United States of America, 110(46), 18454–18459. http://doi.org/10.1073/pnas.1318271110/-/DCSupplemental

Bakuza, J. S., & Nkwengulila, G. (2009). Variation over time in parasite prevalence among free-ranging chimpanzees at Gombe National Park, Tanzania. International Journal of Primatology, 30(1), 43–53. http://doi.org/10.1007/s10764-008-9329-7

Balmer, O., Stearns, S. C., Schötzau, A., & Brun, R. (2009). Intraspecific competition between co-infecting parasite strains enhances host survival in African trypanosomes. Ecology, 90(12), 3367–3378.

Buckner, J. C., Lynch Alfaro, J. W., Rylands, A. B., & Alfaro, M. E. (2015). Biogeography of the marmosets and tamarins (Callitrichidae). Molecular Phylogenetics and Evolution, 82 Pt B, 413–425. http://doi.org/10.1016/j.ympev.2014.04.031

Cardon, M., Loot, G., Grenouillet, G., & Blanchet, S. (2011). Host characteristics and environmental factors differentially drive the burden and pathogenicity of an ectoparasite: a multilevel causal analysis. Journal of Animal Ecology, 80(3), 657–667. http://doi.org/10.1111/j.1365-2656.2011.01804.x

Carrasco, F., Tantaleán, M., Gibson, K. N., & Williams, M. (2008). Prevalence of intestinal helminths from a population of wild spider monkeys *ateles belzebuth chamek* in Manu National Park, Peru. Neotropical Helminthology, 2(1), 19–26.

Chapman, C. A., Gillespie, T. R., & Speirs, M. L. (2005). Parasite prevalence and richness in sympatric colobines: effects of host density. American Journal of Primatology, 67(2), 259–266. http://doi.org/10.1002/ajp.20181

Clough, D., Heistermann, M., & Kappeler, P. M. (2010). Host intrinsic determinants and potential consequences of parasite infection in free-ranging red-fronted lemurs *(Eulemur fulvus rufus)*. American Journal of Physical Anthropology, 142(3), 441–452. http://doi.org/10.1002/ajpa.21243Clutton-

Brock, T., & Sheldon, B. C. (2010). Individuals and populations: the role of long-term, individual-based studies of animals in ecology and evolutionary biology. Trends in Ecology & Evolution, 25(10), 562–573. http://doi.org/10.1016Zj.tree.2010.08.002

Cogswell, F. (2007). Parasites of non-human primates. In D. G. Baker (Ed.), Flynn’s Parasites of Laboratory Animals (2nd ed., pp. 693–743). Ames: John Wiley & Sons.

Collins, W. E., & Jeffery, G. M. (2007). *Plasmodium malariae:* parasite and disease. Clinical Microbiology Reviews, 20(4), 579–592. http://doi.org/10.1128/CMR.00027-07

Conga, D. F., Bowler, M., Tantaleán, M., Montes, D., Serra-Freire, N. M., & Mayor, P. (2013). Intestinal helminths in wild Peruvian red uakari monkeys (*Cacajao calvus ucayalii*)in the northeastern Peruvian Amazon. Journal of Medical Primatology, 43(2), 130–133. http://doi.org/10.1111/jmp.12092

Cosgrove, G. E., Nelson, B., & Gengozian, N. (1968). Helminth parasites of the tamarin, *Saguinus fuscicollis*. Laboratory Animal Care, 18, 654–656.

Cote, I. M., & Poulin, R. (1995). Parasitism and group size in social animals: A meta-analysis. Behavioral Ecology, 6(2), 159–165. http://doi.org/10.1093/beheco/6.2.159

Cox, F. E. (2001). Concomitant infections, parasites and immune responses. Parasitology, 122 Suppl, S23–38.

Erkenswick, G. A., Watsa, M., Gozalo, A. S., Dmytryk, N., & Parker, P. G. (2017a). Temporal and demographic blood parasite dynamics in two free-ranging neotropical primates. International Journal for Parasitology: Parasites and Wildlife, 6(2), 59–68. http://doi.org/10.1016/j.ijppaw.2017.03.004

Erkenswick, G. A., Watsa, M., Pacheco, M. A., Escalante, A. A., & Parker, P. G. (2017b). Chronic *Plasmodium brasilianum* infections in wild Peruvian tamarins. PLoS One, 12(9), e0184504-14. http://doi.org/10.1371/journal.pone.0184504

Freeland, W. (1976). Pathogens and the evolution of primate sociality. Biotropica, 8(1), 12–24.

Freeland, W. J. (1979). Primate social groups as biological islands. Ecology, 60(4), 719. http://doi.org/10.2307/1936609

Gillespie, T. R. (2006). Noninvasive assessment of gastrointestinal parasite infections in free ranging primates. International Journal of Primatology, 27(4), 1129–1143. http://doi.org/10.1007/s10764-006-9064-x

Gillespie, T. R., & Chapman, C. A. (2008). Forest fragmentation, the decline of an endangered primate, and changes in host-parasite interactions relative to an unfragmented forest. American Journal of Primatology, 70(3), 222–230. http://doi.org/10.1002/ajp.20475

Gillespie, T. R., Barelli, C., & Heistermann, M. (2013). Effects of social status and stress on patterns of gastrointestinal parasitism in wild white-handed gibbons *(Hylobates lar)*. American Journal of Physical Anthropology, 150(4), 602–608. http://doi.org/10.1002/ajpa.22232

Gillespie, T. R., Chapman, C. A., & Greiner, E. C. (2005). Effects of logging on gastrointestinal parasite infections and infection risk in African primates. Journal of Applied Ecology, 42(4), 699–707. http://doi.org/10.1111/j.1365-2664.2005.01049.x

Gillespie, T. R., Lonsdorf, E. V., Canfield, E. P., Meyer, D. J., Nadler, Y., Raphael, J., et al. (2010). Demographic and ecological effects on patterns of parasitism in eastern chimpanzees *(Pan troglodytes schweinfurthii)* in Gombe National Park, Tanzania. American Journal of Physical Anthropology, 143(4), 534–544. http://doi.org/10.1002/ajpa.21348

Gordo, M., Calleia, F. O., Vasconcelos, S. A., Leite, J. J. F., & Ferrari, S. F. (2013). The challenges of survival in a concrete jungle: conservation of the pied tamarin *(Saguinus bicolor)* in the urban landscape of Manaus, Brazil. In L. K. Marsh (Ed.), Primates in Fragments (pp. 357–370). New York: Springer New York. http://doi.org/10.1007/978-1-4614-8839-2_23

Guerrero M, F., Serrano-Martínez, E., Tantaleán, M., V, Quispe H, M., & Casas, G., V. (2012). Identificacion de parasitos gastrointestinales en primates no humanos del zoológico parque natural de Pucallpa, Perú. Revista De Investigaciones Veterinarias Del Perú, 23(4), 469–478.

Haukisalmi, V., Henttonen, H., & Tenora, F. (1988). Population dynamics of common and rare helminths in cyclic vole populations. Journal of Animal Ecology, 57, 807–825.

Heymann, E. W., Knogge, C., & Tirado Herrera, E. R. (2000). Vertebrate predation by sympatric tamarins, *Saguinus mystax* and *Saguinus fuscicollis*. American Journal of Primatology, 51(2), 153–158.

Holm, S. (1979). A simple sequentially rejective multiple test procedure. Scandinavian Journal of Statistics, 6, 65–70. http://doi.org/10.2307/4615733

Hopkins, M. E., & Nunn, C. L. (2007). A global gap analysis of infectious agents in wild primates. Diversity and Distributions, 13(5), 561–572. http://doi.org/10.1111/.1472-4642.2007.00364.x

Howells, M. E., Pruetz, J., & Gillespie, T. R. (2011). Patterns of gastro-intestinal parasites and commensals as an index of population and ecosystem health: the case of sympatric western chimpanzees (*Pan troglodytes verus*) and guinea baboons (*Papio hamadryas papio*) at Fongoli, Senegal. American Journal of Primatology, 73(2), 173–179. http://doi.org/10.1002/ajp.20884

Hudson, P. J. (1998). Prevention of Population Cycles by Parasite Removal. Science, 282(5397), 2256–2258. http://doi.org/10.1126/science.282.5397.2256

Hudson, P. J., Dobson, A. P., & Lafferty, K. D. (2006). Is a healthy ecosystem one that is rich in parasites? Trends in Ecology & Evolution, 21(7), 381–385. http://doi.org/10.10167j.tree.2006.04.007

King, N. W., Jr. (1993). Prosthenorchiasis. In Nonhuman Primates (pp. 65–68). Berlin, Heidelberg: Springer Berlin Heidelberg. http://doi.org/10.1007/978-3-642-84924-4_15

Knowles, S. C. L., Fenton, A., Petchey, O. L., Jones, T. R., Barber, R., & Pedersen, A. B. (2013). Stability of within-host-parasite communities in a wild mammal system. Proceedings of the Royal Society of London. Series B, Biological Sciences, 280(1762), 20130598–20130598. http://doi.org/10.1016/j.pt.2005.06.011

Lalremruata, A., Magris, M., Vivas-Martínez, S., Koehler, M., Esen, M., Kempaiah, P., et al. (2015). Natural infection of *Plasmodium brasilianum* in humans: Man and monkey share quartan malaria parasites in the Venezuelan Amazon. EBioMedicine, 2(9), 1186–1192. http://doi.org/10.1016/j.ebiom.2015.07.033

Lee, G. Y., Boyce, W. M., & Orr, K. (1996). Diagnosis and treatment of lungworm *(Filariopsis arator;* metastrongyloidea: filaroididae) infection in white-faced capuchins *(Cebus capucinus)*. Journal of Zoo and Wildlife Medicine, 27(2), 197–200.

Leite, G. C., Duarte, M. H. L., & Young, R. J. (2011). Human-marmoset interactions in a city park. Applied Animal Behaviour Science, 132(3-4), 187–192. http://doi.org/10.1016/j.applanim.2011.03.013

Lello, J., Boag, B., Fenton, A., Stevenson, I. R., & Hudson, P. J. (2004). Competition and mutualism among the gut helminths of a mammalian host. Nature, 428(6985), 840–844. http://doi.org/10.1038/nature02472

MacIntosh, A. J. J., Hernandez, A. D., & Huffman, M. A. (2010). Host age, sex, and reproductive seasonality affect nematode parasitism in wild Japanese macaques. Primates, 51(4), 353–364. http://doi.org/10.1007/s10329-010-0211-9

MacIntosh, A. J. J., Jacobs, A., Garcia, C., Shimizu, K., Mouri, K., Huffman, M. A., & Hernandez, A. D. (2012). Monkeys in the middle: parasite transmission through the social network of a wild primate. PLoS One, 7(12), e51144. http://doi.org/10.1371/journal.pone.0051144.t004

Maizels, R. M., & McSorley, H. J. (2016). Regulation of the host immune system by helminth parasites. Journal of Allergy and Clinical Immunology, 138(3), 666–675. http://doi.org/10.1016/jjaci.2016.07.007

Matauschek, C., Roos, C., & Heymann, E. W. (2011). Mitochondrial phylogeny of tamarins *(Saguinus, Hoffmannsegg* 1807) with taxonomic and biogeographic implications for the *S. nigricollis* species group. American Journal of Physical Anthropology, 144(4), 564–574. http://doi.org/10.1002/ajpa.21445

Michaud, C., Tantaleán, M., Ique, C., Montoya, E., & Gozalo, A. (2003). A survey for helminth parasites in feral New World non-human primate populations and its comparison with parasitological data from man in the region. Journal of Medical Primatology, 32(6), 341–345.

Milton, K. (1996). Effects of bot fly *(Alouattamyia baeri)* parasitism on a free-ranging howler monkey *(Alouatta palliata)* population in Panama. Journal of Zoology, 239, 39–63.

Monteiro, R. V., Dietz, J. M., Beck, B. B., Baker, A. J., Martins, A., & Jansen, A. M. (2007a). Prevalence and intensity of intestinal helminths found in free-ranging golden lion tamarins *(Leontopithecus rosalia*, Primates, Callitrichidae) from Brazilian Atlantic forest. Veterinary Parasitology, 145(1-2), 77–85. http://doi.org/10.1016/j.vetpar.2006.12.004

Monteiro, R. V., Dietz, J. M., Raboy, B., Beck, B., Vleeschower, K. D., Baker, A., et al. (2007b). Parasite community interactions: *Trypanosoma cruzi* and intestinal helminths infecting wild golden lion tamarins *Leontopithecus rosalia* and golden-headed lion tamarins *L. chrysomelas* (Callitrichidae, L., 1766). Parasitology Research, 101(6), 1689–1698. http://doi.org/10.1007/s00436-007-0652-2

Muehlenbein, M. P., & Watts, D. P. (2010). The costs of dominance: testosterone, cortisol and intestinal parasites in wild male chimpanzees. BioPsychoSocial Medicine, 4(1), 21. http://doi.org/10.1186/1751-0759-4-21

Muller, B. (2007). Determinants of the diversity of intestinal parasite communities in sympatric New World primates (*Saguinus mystax, Saguinus fuscicollis, Callicebus cupreus*). Tierärztliche Hochschule Hannover.

Nunn, C. L., & Altizer, S. M. (2005). Global Mammal Parasite Database: an online resource for infectious disease records in wild primates. Retrieved April 29, 2013, from http://www.mammalparasites.org

Nunn, C. L., & Heymann, E. W. (2005). Malaria infection and host behavior: a comparative study of Neotropical primates. Behavioral Ecology and Sociobiology, 59(1), 30–37. http://doi.org/10.1007/s00265-005-0005-z

Nunn, C. L., Altizer, S., Jones, K. E., & Sechrest, W. (2003). Comparative tests of parasite species richness in primates. The American Naturalist, 162(5), 597–614. http://doi.org/10.1086/378721

Nunn, C. L., Brezine, C., Jolles, A. E., & Ezenwa, V. O. (2014). Interactions between micro-and macroparasites predict microparasite species richness across primates. The American Naturalist, 183(4), 1–14. http://doi.org/10.5061/dryad.q749f

Orkin, J. D., Yang, Y., Yang, C., Yu, D. W., & Jiang, X. (2016). Cost-effective scat-detection dogs: unleashing a powerful new tool for international mammalian conservation biology. Scientific Reports, 6, 34758. http://doi.org/10.1038/srep34758

Parr, N. A., Fedigan, L. M., & Kutz, S. J. (2013). A Coprological Survey of Parasites in White-Faced Capuchins *(Cebus capucinus)* from Sector Santa Rosa, ACG, Costa Rica. Folia Primatologica, 84(2), 102–114. http://doi.org/10.1159/000348287

Petney, T. N., & Andrews, R. H. (1998). Multiparasite communities in animals and humans: frequency, structure and pathogenic significance. International Journal for Parasitology: Parasites and Wildlife, 28(3), 377–393.

Phillips, K. A., Haas, M. E., Grafton, B. W., & Yrivarren, M. (2004). Survey of the gastrointestinal parasites of the primate community at Tambopata National Reserve, Peru. Journal of Zoology, 264(2), 149–151. http://doi.org/10.1017/S0952836904005680

Pontes, A. R. M., & da Cruz, M. A. O. M. (1995). Home range, intergroup transfers, and reproductive status of common marmosets *Callithrix jacchus* in a forest fragment in NorthEastern Brazil. Primates, 36(3), 335–347. http://doi.org/10.1007/BF02382857

Porter, J. A. (1972). Parasites of marmosets. Laboratory Animal Science, 22(4), 503–506.

R Development Core Team. (2015). R: A language and Environment for Statistical Computing. Vienna, Austria. Retrieved from https://www.R-project.org/

Rifkin, J. L., Nunn, C. L., & Garamszegi, L. Z. (2012). Do animals living in larger groups experience greater parasitism? A meta-analysis. The American Naturalist, 180(1), 70–82. http://doi.org/10.1086/666081

Rosin, C., & Swamy, V. (2013). Variable density responses of primate communities to hunting pressure in a western Amazonian river basin. Neotropical Primates, 20(1), 25–31. http://doi.org/10.1896/044.020.0105

Rylands, A. B., & Mittermeier, R. A. (2008). Saguinus imperator. In: IUCN 2012. IUCN Red List of Threatened Species. Version 2012.2. Retrieved April 10, 2013, from

Salzer, J. S., Deutsch, J. C., Raño, M., Kuhlenschmidt, M. S., & Gillespie, T. R. (2010). Black and gold howler monkeys *(Alouatta caraya)* as sentinels of ecosystem health: patterns of zoonotic protozoa infection relative to degree of human-primate contact. American Journal of Primatology, 73(1), 75–83. http://doi.org/10.1002/ajp.20803

Setchell, J. M., Charpentier, M. J. E., Bedjabaga, I.-B., Reed, P., Wickings, E. J., & Knapp, L. A. (2006). Secondary sexual characters and female quality in primates. Behavioral Ecology and Sociobiology, 61(2), 305–315. http://doi.org/10.1007/s00265-006-0260-7

Setchell, J. M., Charpentier, M., & Abbott, K. M. (2009). Is brightest best? Testing the Hamilton-Zuk hypothesis in mandrills. International Journal of Primatology, 30, 825–844. http://doi.org/10.1007/s10764-009-9371-0

Sikes, R. S., & Gannon, W. L. (2011). Guidelines of the American Society of Mammalogists for the use of wild mammals in research. Journal of Mammalogy, 92(1), 235–253. http://doi.org/10.1644/10-MAMM-F-355.1

Smith, A. C. (2000). Interspecific differences in prey captured by associating saddleback *(Saguinus fuscicollis)* and moustached *(Saguinus mystax)* tamarins. Journal of Zoology, 251(3), 315–324. http://doi.org/10.1111/j.1469-7998.2000.tb01082.x

Soto-Calderón, I. D., Acevedo-Garcés, Y. Á., Alvarez-Cardona, J., Hernñndez-Castro, C., & García-Montoya, G. M. (2016). Physiological and parasitological implications of living in a city: the case of the white-footed tamarin *(Saguinus leucopus)*. American Journal of Primatology, 78(12), 1272–1281. http://doi.org/10.1002/ajp.22581

Steinfelder, S., O’Regan, N. L., & Hartmann, S. (2016). Diplomatic Assistance: Can Helminth- Modulated Macrophages Act as Treatment for Inflammatory Disease? PLoS Pathogens, 12(4), e1005480-14. http://doi.org/10.1371/journal.ppat.1005480

Strait, K., Else, J. G., & Eberhard, M. L. (2012). Parasitic diseases of nonhuman primates. In Nonhuman primates in biomedical research (pp. 197–297). Elsevier. http://doi.org/10.1016/B978-0-12-381366-4.00004-3

Stuart, M., Pendergast, V., Rumfelt, S., Pierberg, S., Greenspan, L., Glander, K., & Clarke, M. (1998). Parasites of Wild Howlers *(Alouatta* spp.). International Journal of Primatology, 19(3), 493–512. http://doi.org/10.1023/A:1020312506375

Sussman, R. W., & Kinzey, W. G. (1984). The ecological role of the Callitrichidae: a review. American Journal of Physical Anthropology, 64(4), 419–449. http://doi.org/10.1002/ajpa.1330640407

Tantaleán, M., Gozalo, A., & Montoya, E. (1990). Notes on some helminth parasites from Peruvian monkeys. Lab Primate Newsletter, 29, 6–8.

Tavela, A. de O., Fuzessy, L. F., e Silva, V. H. D., da Silva, F. de F. R., Junior, M. C., Silva, I. de O., & Souza, V. B. (2013). Helminths of wild hybrid marmosets *(Callithrix* sp.) living in an environment with high human activity. Revista Brasileira De Parasitologia Veterinária, 22(3), 391–397. http://doi.org/10.1590/S1984-29612013000300012

Telfer, S., Birtles, R., Bennett, M., Lambin, X., Paterson, S., & Begon, M. (2008). Parasite interactions in natural populations: insights from longitudinal data. Parasitology, 135(7), 767–781. http://doi.org/10.1017/S0031182008000395

Toft, D. J., & Eberhard, M. L. (1998). Parasitic diseases. In T. B. Bennet, C. R. Abee, & R. Henrickson (Eds.), Nonhuman Primates in Biomedical Research (pp. 111–205). San Diego: Elsevier Inc.

Valdespino, C., Rico-Hernñndez, G., & Mandujano, S. (2010). Gastrointestinal parasites of Howler monkeys *(Alouatta palliata)* inhabiting the fragmented landscape of the Santa Marta mountain range, Veracruz, Mexico. American Journal of Primatology, 72, 539–548. http://doi.org/10.1002/ajp.20808

Vitone, N. D., Altizer, S., & Nunn, C. L. (2004). Body size, diet and sociality influence the species richness of parasitic worms in anthropoid primates. Evolutionary Ecology, 6(2), 183199.

Walker, S. F., Bosch, J., Gomez, V., Garner, T. W. J., Cunningham, A. A., Schmeller, D. S., et al. (2010). Factors driving pathogenicity vs. prevalence of amphibian panzootic chytridiomycosis in Iberia. Ecology Letters, 13(3), 372–382. http://doi.org/10.1111/j.1461-0248.2009.01434.x

Walther, B. A., Cotgreave, P., Price, R. D., & Gregory, R. D. (1995). Sampling effort and parasite species richness. Parasitology Today, 11(8), 306–310. http://doi.org/10.1016/0169-4758(95)80047-6

Watsa, M. (2013). Growing up tamarin: morphology, reproduction, and population demography of sympatric free-ranging *Saguinus fuscicollis* and *S. imperator*. Washington University in St. Louis.

Watsa, M. (2015, December 11). 200,000 of Peru’s primates trafficked for pet trade or bushmeat yearly. Retrieved March 14, 2017, from https://news.mongabay.com/2015/12/200000-of-perus-primates-trafficked-for-pet-trade-or-bushmeat-yearly/

Watsa, M., Erkenswick, G. A., Rehg, J. A., & Pitman, R. L. (2012). Distribution and new sightings of Goeldi’s monkey *(Callimico goeldii)* in Amazonian Perú. International Journal of Primatology, 33(6), 1477–1502. http://doi.org/10.1007/s10764-012-9632-1

Watsa, M., Erkenswick, G., & Robakis, E. (2017). Modeling developmental class provides insights into individual contributions to infant survival in callitrichids. International Journal of Primatology, 38(6), 1032–1057. http://doi.org/10.1007/s10764-017-9995-4

Watsa, M., Erkenswick, G., Halloran, D., Kane, E. E., Poirier, A., Klonoski, K., et al. (2015). A field protocol for the capture and release of callitrichids. Neotropical Primates, 22, 59–68.

Wenz, A., Heymann, E. W., Petney, T. N., & Taraschewski, H. F. (2009). The influence of human settlements on the parasite community in two species of Peruvian tamarin. Parasitology, 137(04), 675. http://doi.org/10.1017/S0031182009991570

Williamson, E. A., & Feistner, A. (2011). Habituating primates: processes, techniques, variables and ethics. In J. M. Setchell & D. J. Curtis (Eds.), Field and laboratory methods in primatology: a practical guide (pp. 33–50). Cambridge, UK: Cambridge University Press.

Wislocki, G. B. (1939). Observations on twinning in marmosets. American Journal of Anatomy, 64(3), 445–483. http://doi.org/10.1002/aja.1000640305

Wolff, P. L. (1990). The parasites of New World primates: a review. Proceedings of the American Association of Zoo Veterinarians, 87–94.

Zajac, A. M., & Conboy, G. A. (2012). Veterinary Clinical Parasitology (8th ed.) (p. 368). Ames, IA: Wiley-Blackwell Publishing.

